# Ad26.COV2.S elicited neutralizing activity against Delta and other SARS-CoV-2 variants of concern

**DOI:** 10.1101/2021.07.01.450707

**Authors:** Mandy Jongeneelen, Krisztian Kaszas, Daniel Veldman, Jeroen Huizingh, Remko van der Vlugt, Theo Schouten, David Zuijdgeest, Taco Uil, Griet van Roey, Nuria Guimera, Marjon Navis, Rinke Bos, Mathieu le Gars, Jerald Sadoff, Leacky Muchene, Jarek Juraszek, Johannes PM Langedijk, Ronald Vogels, Jerome Custers, Hanneke Schuitemaker, Boerries Brandenburg

## Abstract

Severe acute respiratory syndrome coronavirus 2 (SARS-CoV-2) continues to evolve and recently emerging variants with substitutions in the Spike protein have led to growing concerns over increased transmissibility and decreased vaccine coverage due to immune evasion. Here, sera from recipients of a single dose of our Ad26.COV2.S COVID-19 vaccine were tested for neutralizing activity against several SARS-CoV-2 variants of concern. All tested variants demonstrated susceptibility to Ad26.COV2.S-induced serum neutralization albeit mainly reduced as compared to the B.1 strain. Most pronounced reduction was observed for the B.1.351 (Beta; 3.6-fold) and P.1 (Gamma; 3.4-fold) variants that contain similar mutations in the receptor-binding domain (RBD) while only a 1.6-fold reduction was observed for the widely spreading B.1.617.2 (Delta) variant.

## Introduction

In response to the emergence of SARS-CoV-2 in 2019, Janssen Vaccines developed the Ad26.COV2.S COVID-19 vaccine. The vaccine is currently being tested in multiple clinical trials and interim immunogenicity and efficacy data have been reported ^1-3^. These showed high responder rates for both humoral and cellular immunity against the original SARS-CoV-2 variants (WA1/2020, Wuhan-Hu-1), and high vaccine efficacy (>80%) against severe COVID-19, with complete protection against COVID-19 related hospitalization and death across geographies, including South Africa, where >95% of cases from whom sequence information was available, were infected with the Beta variant of concern (VOC) of SARS-CoV-2 (B.1.351 lineage) ^1^.

The single dose Ad26.COV2.S COVID-19 vaccine was granted emergency use or (conditional) marketing authorization in more than 50 countries and as of July 2021, more than 19 million people have been vaccinated worldwide.

We recently demonstrated that the neutralizing activity in serum against the Beta (B.1.351) and Gamma (P.1) variants were respectively 5.0 and 3.3-fold reduced as compared to the WA1/2020 variant. However, non-neutralizing Fc mediated antibody functions such as antibody dependent cellular phagocytosis, complement deposition and NK cell activation as well as CD4 and CD8 T cell responses, including central memory and effector memory responses, were comparable across WA1/2020, B.1.1.7, B.1.351, P.1 and CAL.20C variants ^4^. The Delta VOC (B.1.617.2), first detected in India, is now rapidly spreading across the globe and information on activity of vaccine elicited immune responses against this variant is urgently needed.

Here we provide an update on the in vitro binding and neutralization titers detected in sera of older adults immunized with Ad26.COV2.S during the phase 3 ENSEMBLE trial (VAC31518COV3001) against the VOCs, including the Delta variant B.1.617.2.

## Methods

### Clinical Trial

Participants of phase 3 ENSEMBLE trial (VAC31518COV3001, ClinicalTrials.gov number, NCT04505722) were immunized with one dose of 5×10^10^ viral particles of replication-incompetent Ad26 vector encoding for the prefusion-stabilized SARS-COV-2 spike protein sequence (Wuhan-Hu-1; GenBank accession no. MN908947). Sera were collected 71 days later (day 71). Study protocols and results have been reported previously ^1^.

### ELISA

IgG binding to SARS-CoV-2 spike protein was measured by ELISA using a recombinant and stabilized trimeric spike protein antigen based on the Wuhan-Hu-1 SARS-CoV-2 strain (GenBank accession no. MN908947, D614) ^5^. The SARS-CoV-2 spike protein antigen (2.0µg/mL) was directly adsorbed on half-area 96-well OptiPlate (96W HB, Perkin Elmer) microplates for 2h at 37°C in a humidified incubator.

Following incubation, plates were washed three times in PBS/0.05% Tween-20 (PBS-T) and blocked with 1% Casein in PBS for 1h at room temperature. Plates were washed with PBS-T after the blocking step, then serum standards (high titer human convalescent and naïve reference sera), control antibodies, and serum samples were serial diluted (2.5-fold) before incubated on the plates for 1h at room temperature. Next, the plates were washed three times with PBS-T then incubated with peroxidase-conjugated Goat anti-Human IgG (Jackson ImmunoResearch) diluted in blocking buffer for 1h at room temperature, washed three times in PBS-T, and developed with detection substrate (Clarity Western ECL peroxide reagent and luminol enhancer, Bio-Rad) for 10min at room temperature and protected from light. Signal was read out on an Envision plate reader (Perkin Elmer) as relative luminescence units (RLUs). Titers are reported as log10 relative potency (log10 RP) compared with a high titer serum sample used as a reference standard, with a lower limit of quantification at 1.218.

### Recombinant lentivirus-based pseudotyped virus neutralization assay (psVNA)

For measuring the breadth of neutralization against tested SARS-CoV-2 spike variants, SARS-CoV-2 spike-neutralizing antibody titers were measured in psVNA against several SARS-CoV-2 spike variants as described previously ^6^. In brief, for the generation of pseudotyped HIV-based lentiviruses, codon optimized and synthesized DNA encoding SARS-CoV-2 Spike protein (based on Wuhan-Hu-1; GenBank accession no. MN908947) C-terminally truncated by 19 amino acids was cloned into a derivative of the pCDNA3.1 expression vector (Thermo Fisher Scientific). Substitutions and deletions in the Spike protein gene open reading frame (shown in **Figure 1**) were introduced using standard molecular biology techniques and confirmed by sequencing. HIV-based lentiviral pseudotyped particles harboring the SARS-CoV-2 Spike protein variants were produced using the ViraPower Lentiviral Expression system (Thermo Fisher Scientific) according to manufacturer’s protocol with minor changes. Neutralization assays for the different variants were performed in parallel on Hek293T target cells stably expressing the human ACE2 and human TMPRSS2 genes (VectorBuilder; Cat. CL0015). Cells were pre-seeded in white half-area 96-well tissue culture plates (Perkin Elmer) at a density of 1.5 × 10^4^ cells/well. Heat-inactivated serum samples were two-fold serial diluted in duplicates over 10 columns in phenol red free DMEM supplemented with 1% FBS and 1% PenStrep. Serum standards (high titer human convalescent and naïve reference sera), controls (including antibodies), and serial diluted serum samples were incubated at room temperature with an equal volume of pseudovirus particles with titers of ∼1 × 10^5^ RLUs luciferase activity. After 1h incubation, the serum-particle mixture was inoculated onto Hek293T.ACE2.TMPRSS2 cells. Luciferase activity was measured 40h after transduction by adding an equal volume of NeoLite substrate (Perkin Elmer) to the wells according to the manufacturer’s protocol, followed by readout of RLUs on the EnSight Multimode Plate Reader (Perkin Elmer). SARS-CoV-2 neutralizing titers were calculated in R using a four-parameter curve fit as the sample dilution at which a 50% reduction (IC50) of luciferase readout was observed compared with luciferase readout in the absence of serum (“High Control”). The starting serum sample dilution of 20 was fixed as the limit of detection (LOD).

**Figure 1:**
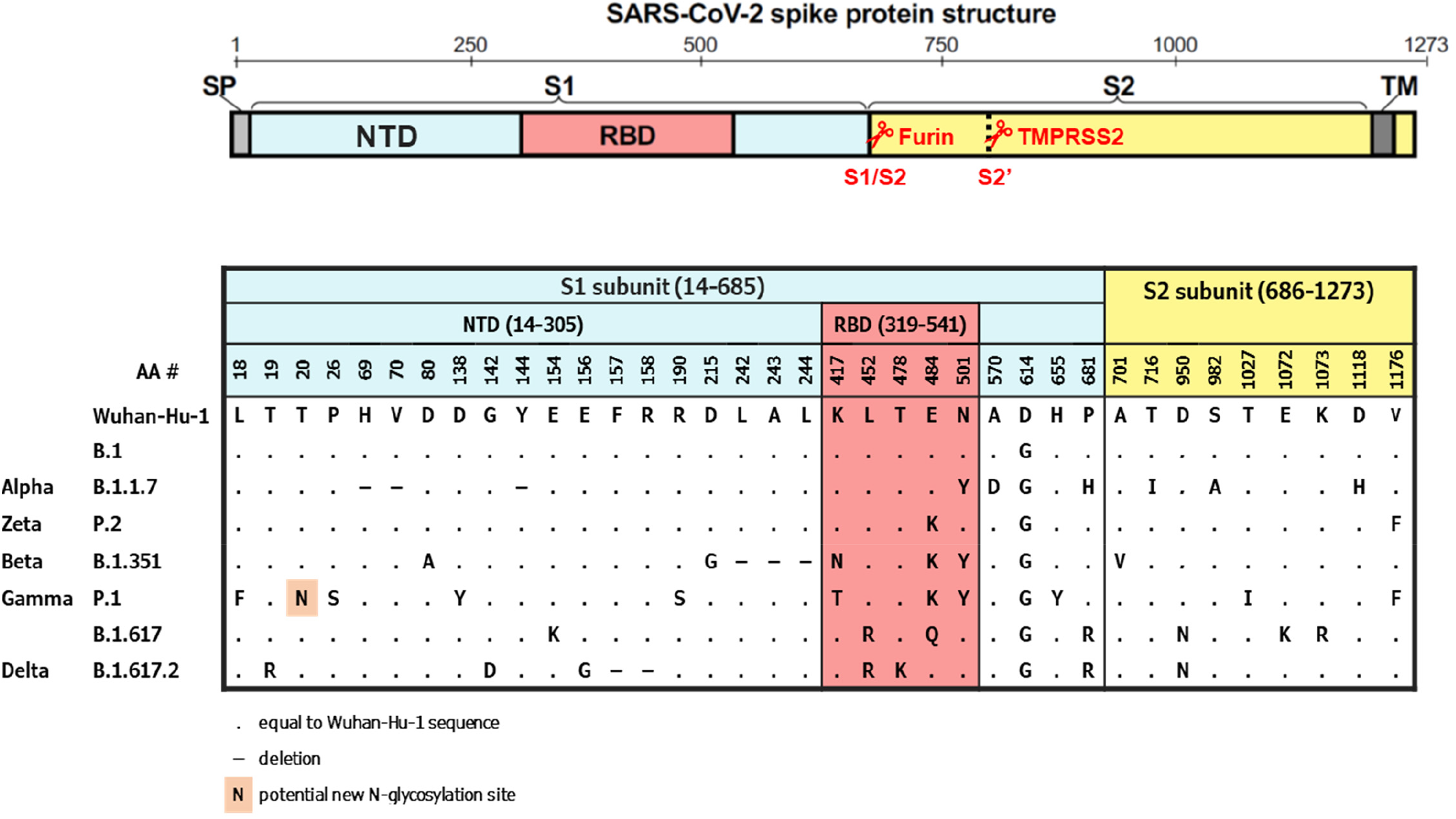
Spike protein schematic structure and alignment of substitutions in SARS-CoV-2 variants evaluated in this study. S, Spike protein; NTD, N-terminal domain; RBD, receptor binding domain; SP, signaling peptide; TM, transmembrane domain; AA, amino-acid.

### Statistical analysis

Geometric mean titers, limit of detection, and fold change to B.1 were reported. Statistical comparison of the fold differences relative to B.1 were performed in SAS 9.4 using a paired-sample T-test at a 5% significance level. No adjustment for multiple comparisons were done. The corresponding 95% confidence intervals were also reported.

## Results

Sera from phase 3 ENSEMBLE trial participants (n=8; from Brazil, RSA, and USA, 71 days post single dose vaccination, age ranging from 47 to 91 years) were evaluated for their ability to bind to B.1 Spike protein in ELISA and for their neutralizing activity against B.1 and variants of concern (listed in Figure 1) in a pseudovirus neutralization assay. Sera were selected based on titer. Binding to B.1 Spike and neutralization titers on B.1 correlated well (Spearman rank correlation coefficient of 0.92, **Figure 2**). Geometric mean neutralization titers (GMT) were observed against all variants. The neutralization titers against B.1.1.7 and B.1 were similar and reduced against all other variants tested, ranging from 1.5 (B.1.617) to 3.6 (B.1.351) fold (**Figure 3**). The Beta (B.1.351) and Gamma (P.1) variants, with mutations at positions 417, 484 and 501 in the receptor-binding domain (RBD), showed the largest reduction in neutralization sensitivity (3.6-fold and 3.4-fold respectively). The Delta (B.1.617.2) variant, with mutations at positions 452 and 478 in the RBD, demonstrated only a 1.6-fold reduction in neutralization sensitivity. Four out of eight individuals had pre-existing Spike antibodies at baseline, as measured by ELISA, 2 of them also tested positive in the psVNA against B.1 at baseline (**Figure 2**). However, the relative neutralizing titers for the Beta (B.1) and Delta (B.1.617.2) variants were similar on day 71 in subjects that were seronegative at baseline compared to the ones with pre-existing ELISA responses at baseline (**Figure 3**).

**Figure 2:**
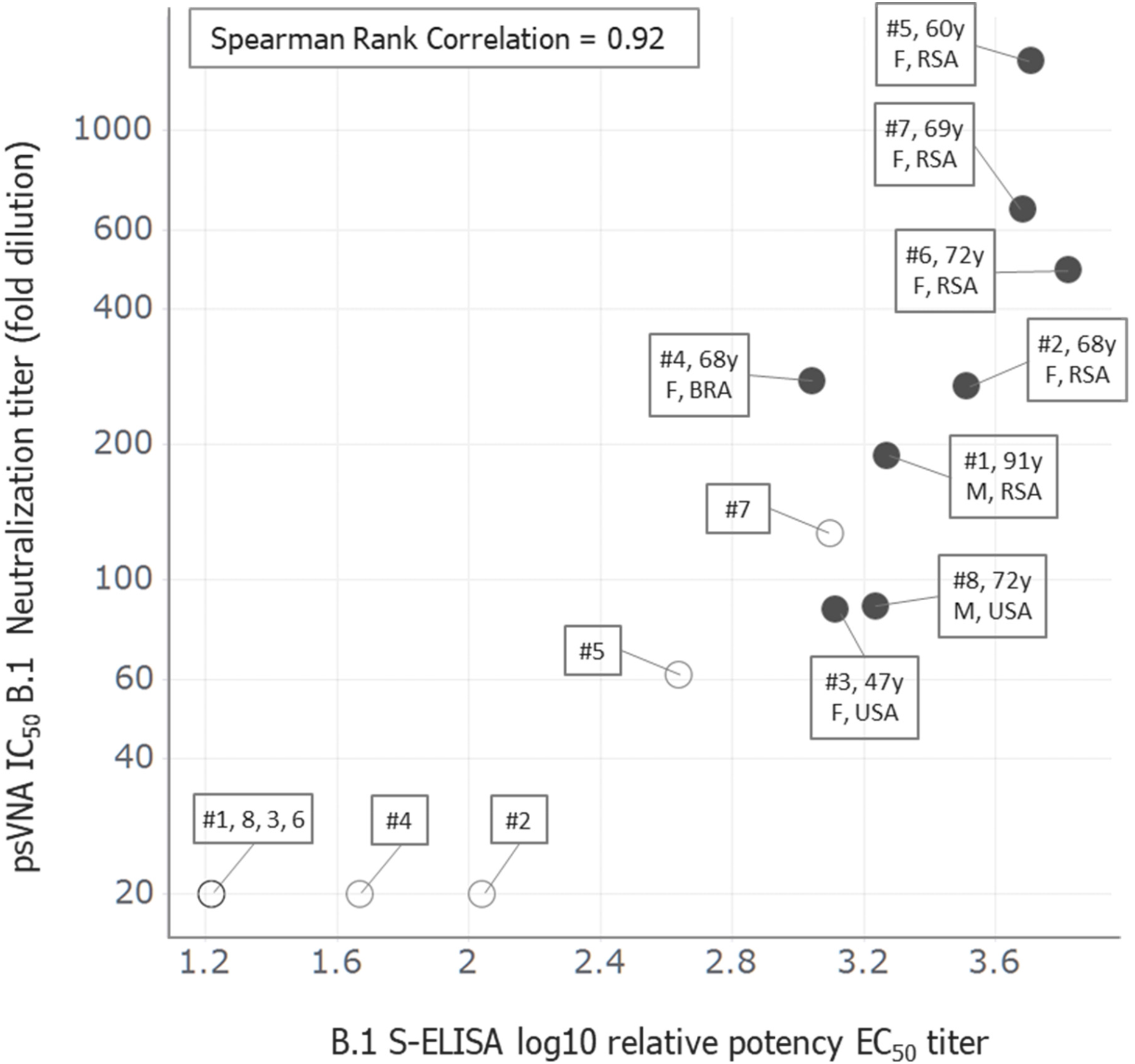
Correlation between SARS-CoV-2 B.1 antibody neutralization and binding titers. Serum samples (N=8) from participants of phase 3 ENSEMBLE trial immunized with one dose of 5×10^10^ viral particles of Ad26.COV2.S were obtained on day 1 (open circle) and day 71 (closed circle) past immunization. Samples were tested in a pseudotyped lentivirus neutralization assay on SARS-CoV-2 B.1 (D614G) and a corresponding B.1 (D614) direct coat S-ELISA. The reciprocal neutralizing titers on the pseudovirus neutralization assay at a 50% inhibitory concentration (IC, serum dilution) versus the log10 relative potency binding titer are shown and each data point represents results from individual serum samples. The additional boxed information next to the data points includes participant number (#), age in years (y), gender (F/M), and country of sampling (RSA, BRA, USA).

**Figure 3:**
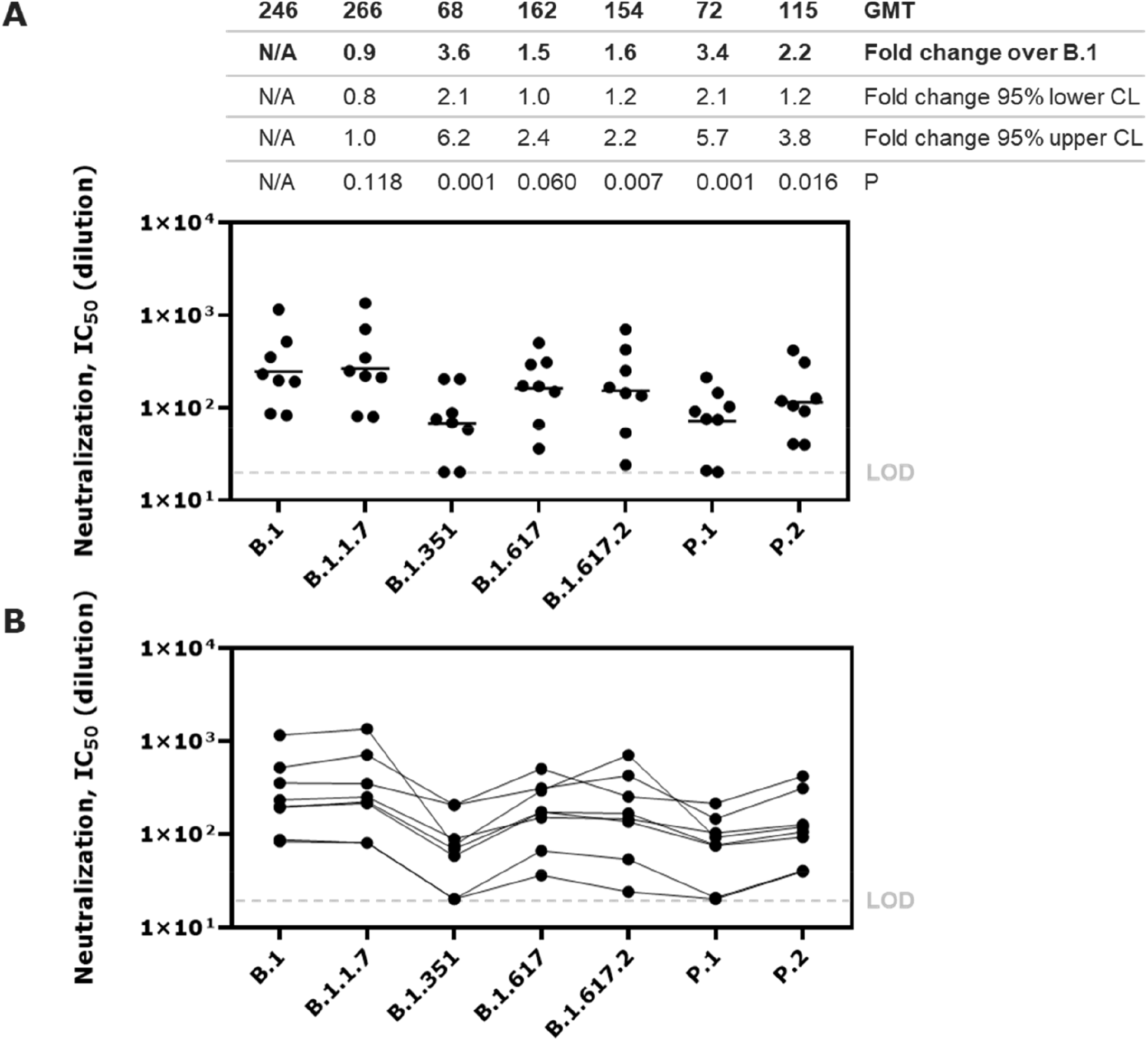
Neutralization activity of serum antibodies elicited by Ad26.COV2.S against SARS-CoV-2 spike pseudotyped virus variants. Serum samples (N=8) from participants of phase 3 ENSEMBLE trial immunized with one dose of 5×10^10^ viral particles of Ad26.COV2.S were obtained on day 71 past immunization. Samples were tested in a pseudotyped lentivirus neutralization assay on SARS-CoV-2 spike protein variants of concern (see Figure 1). The neutralizing titers on the pseudovirus neutralization assay at a 50% inhibitory concentration (IC, serum dilution) are shown. Dots represent results from individual serum samples. A) Shows independent measurements per virus variant, geometric mean titers (GMT) as number and horizontal line, fold changes compared to the B.1, the 95% lower and upper confidence level (CL), as well as the P value. B) Shows the same data with lines connecting matched serum samples.

We recently reported that cellular immunity is conserved against SARS-CoV-2 variants of concern ^4^. While we have not yet tested this for the Delta VOC, in silico analysis has demonstrated the preservation of dominant T cell epitopes in this variant (data not shown).

## Discussion

As compared to the neutralizing activity in Ad26.COV2.S elicited immune sera against the B.1 virus, neutralizing activity is more strongly reduced against the Beta (B.1.351) and Gamma (P.1) variants than against the rapidly spreading Delta (B.1.617.2) variant. These results are in line with recently published studies in which sera from subjects who received the Moderna, Pfizer-BioNTech, or Oxford-AstraZeneca COVID-19 vaccines were tested for neutralizing activity against VOCs ^7, 8^. For all the vaccines, the reduction in neutralization titer was greater for the Beta (B.1.351) than observed for the Delta (B.1.617.2) variant.

At this point, no vaccine efficacy against the Delta variant is available although Real World Evidence studies have suggested that Pfizer-BioNTech and Astra-Zeneca vaccines are effective against this new variant ^9^. The Janssen COVID-19 vaccine efficacy (VE) against the Delta variant of concern is currently unknown and may become available from our ongoing phase 3 trials only later this year. However, we have observed high VE against severe COVID-19 in South Africa, with full protection against COVID-19 related hospitalization and death ^1^, while >95% of cases with available sequence information were classified as the Beta (B.1.351) variant against which neutralizing activity on day 29 was more severely impacted. This strongly suggests that VE of a single dose of Ad26.COV2.S against Delta variant will be preserved as well, either because lower neutralizing antibody titers are still sufficient to protect or by the contribution of non-neutralizing antibody functions and the strong cellular immune responses that Ad26.COV2.S elicits.

## Contributions

Conceptualization: B.B., M.J., K.K, G.R., N.G., M.G, J.S., J.C., H.S.; methodology: M.J., D.V., J.H., R.vdV., T.S., D.Z., T.U.; formal & statistical analysis: M.J., M.N., R.B., K.K., L.M., J.J.; writing, review, and editing: B.B., H.S., M.J., K.K, R.V., N.G., J.C. All authors have read and agreed to the published version of the manuscript.

## Acknowledgments

We thank all participants of our clinical trials. We thank Lucy Rutten for providing the S-ELISA antigen.

